# Determinants of high residual post-PCV13 pneumococcal vaccine type carriage in Blantyre, Malawi: a modelling study

**DOI:** 10.1101/477695

**Authors:** J. Lourenço, U. Obolski, T.D. Swarthout, A. Gori, N. Bar-Zeev, D. Everett, A.W. Kamng’ona, T.S. Mwalukomo, A.A. Mataya, C. Mwansambo, M. Banda, S. Gupta, N. French, R.S. Heyderman

## Abstract

**Background:** In November 2011, Malawi introduced the 13-valent pneumococcal conjugate vaccine (PCV13) into the routine infant schedule. Four to seven years after introduction (2015-2018), rolling prospective nasopharyngeal carriage surveys were performed in the city of Blantyre. Carriage of *Streptococcus pneumoniae* vaccine serotypes (VT) remained higher than reported in developed countries, and VT impact was surprisingly asymmetric across age-groups. A dynamic transmission model was fit to survey data using a Bayesian Markov-chain Monte Carlo approach, to obtain insights into the determinants of post-PCV13 age-specific VT carriage.

**Results:** Accumulation of naturally acquired immunity with age and age-specific transmission potential were both key to reproducing the observed data. VT carriage reduction peaked sequentially over time, earlier in younger and later in older age-groups. Estimated vaccine efficacy (protection against carriage) was 66.87% (95% CI 50.49-82.26%), similar to previous estimates. Ten-year projected vaccine impact (VT carriage reduction) among 0-9 years old was lower than observed in other settings, at 76.23% (CI 95% 68.02-81.96%), with sensitivity analyses demonstrating this to be mainly driven by a high local force of infection.

**Conclusions:** We have identified both vaccine-related and host-related determinants of post-PCV13 pneumococcal VT transmission in Blantyre with vaccine impact determined by age-related characteristics of the local force of infection. These findings are likely to be generalisable to other Sub-Saharan African countries in which PCV impact has been lower than desired, and have implications for the interpretation of post-PCV carriage studies and future vaccination programs.

## Introduction

*Streptococcus pneumoniae* (pneumococcus) is a bacterial human pathogen commonly carried asymptomatically in the nasopharynx, which in a minority of carriers can cause severe disease such as pneumonia, meningitis or bacteremia^1^, posing a serious mortality risk, especially for young children (<5 years of age), the elderly (>65 years of age) and the immunocompromised^2^. Pneumococcal carriage is a necessary precursor of severe disease^3^ and transmission, such that reduction of carriage through active control is an important, universal public health goal.

Currently, pneumococcal conjugate vaccines (PCV) are the best available tool to reduce carriage and disease both within risk groups and the general population. These vaccines have consisted of either 7, 10 or 13 polysaccharides conjugated to a carrier protein (PCV7, PCV10, PCV13, respectively). All have been demonstrated to be highly protective against 7, 10 or 13 common pneumococcal serotypes associated with carriage and disease (also termed vaccine serotypes, VT). A frequently observed consequence of PCV introduction is the increase in both carriage and disease of non-VT pneumococci (NVT), likely due to increased niche availability and reduction of competition between VT and NVT^4–9^.

PCV routine vaccination has been a common control strategy for over a decade in developed countries, with past experience showing that both pre- and post-PCV pneumococcal carriage can be highly variable within and between countries^10–16^. PCV vaccines have only recently been introduced in Sub-Saharan African countries, such as Kenya^17,18^, Malawi^19^, The Gambia^20^ and South Africa^21^. In November 2011, Malawi introduced the 13-valent pneumococcal conjugate vaccine (PCV13) as part of the national extended program of immunization with a 3+0 schedule (at 6, 10 and 14 weeks of age). With high routine coverage (~90%) and a small catch-up campaign of young children, PCV13 was expected to quickly reduce carriage as previously reported in developed countries. However, recently published data on nasopharyngeal carriage as measured in a cross-sectional observational study in Blantyre (Southern Malawi), four to seven years after PCV13 introduction (2015-2018), has shown that vaccine impact (VT carriage reduction) has been slower than expected and heterogeneous across age-groups^22^. Epidemiological mathematical models have previously been employed successfully to improve our understanding of pneumococcal dynamics^5,9,23–27^, as well as having contributed to explain, estimate and project PCV impact^8,11,28^. The main advantage of models is their cost-free potential to test hypotheses and gain a mechanistic, ecological and immunological understanding of carriage and disease dynamics, estimating epidemiological parameters which are difficult to otherwise quantify from raw epidemiological data. For example, models have successfully yielded estimates of VT and non-VT pneumococci transmission potentials^26,29–31^, pneumococcal competition factors^8,9,23,28,32,33^ and measures of vaccine-induced protection from carriage at the individual level^11,17,28,34,35^, none of which are readily observed or quantified in cross-sectional observational studies.

In this study we use a Bayesian Markov chain Monte Carlo fitting approach and a dynamic model to investigate the post-PCV13 pneumococcal VT carriage dynamics in Blantyre, Malawi. We find that natural immunity and age-specific transmission potentials are necessary to reproduce observed VT carriage. When compared to numerous literature reports from other regions, our estimated vaccine efficacy (individual-level protection from carriage) was close to expected values, but impact (population-level reduction of VT carriage) was lower both in the short- and long-term. We show that vaccine impact was likely being offset by a high local force of infection compared to other regions of the world. Our study offers key insights into the lower than expected PCV13 impact in Malawi and more generally on the heterogeneous nature of pre- and post-vaccination pneumococcal VT carriage across age-groups and regions. These results can be translated to other Sub-Saharan African countries in which PCV impact has been lower than desired.

## Methods

### Prospective cross-sectional observational study

An observational study using stratified random sampling was conducted to measure pneumococcal nasopharyngeal carriage in Blantyre, Malawi^22^. Sampling was performed twice a year, between June and August 2015 (survey 1), October 2015 and April 2016 (survey 2), May and October 2016 (survey 3), November 2016 and April 2017 (survey 4), May and October 2017 (survey 5); November 2017 and June 2018 (survey 6), and June and December 2018 (survey 7). In this study, we use the mid-point dates of the surveys for model fitting and presentation of results. A total of 7148 individuals were screened with nasopharyngeal swabs processed following WHO recommendations^36^. Isolates were serotyped by latex agglutination (ImmuLex™ 7-10-13-valent Pneumotest; Statens Serum Institute, Denmark). In this study, we use all the data from three age-groups: 499 vaccinated children 2 years old, 2565 vaccinated children 3–7 years old and 1402 unvaccinated children 3–10 years old. For the first three surveys, data on vaccinated 2 years old individuals was not collected. Observed VT carriage levels are presented in Figure 1d (and Table S7). Further details on collection, processing and observations, have been previously described in detail ^22^.

**Figure 1:**
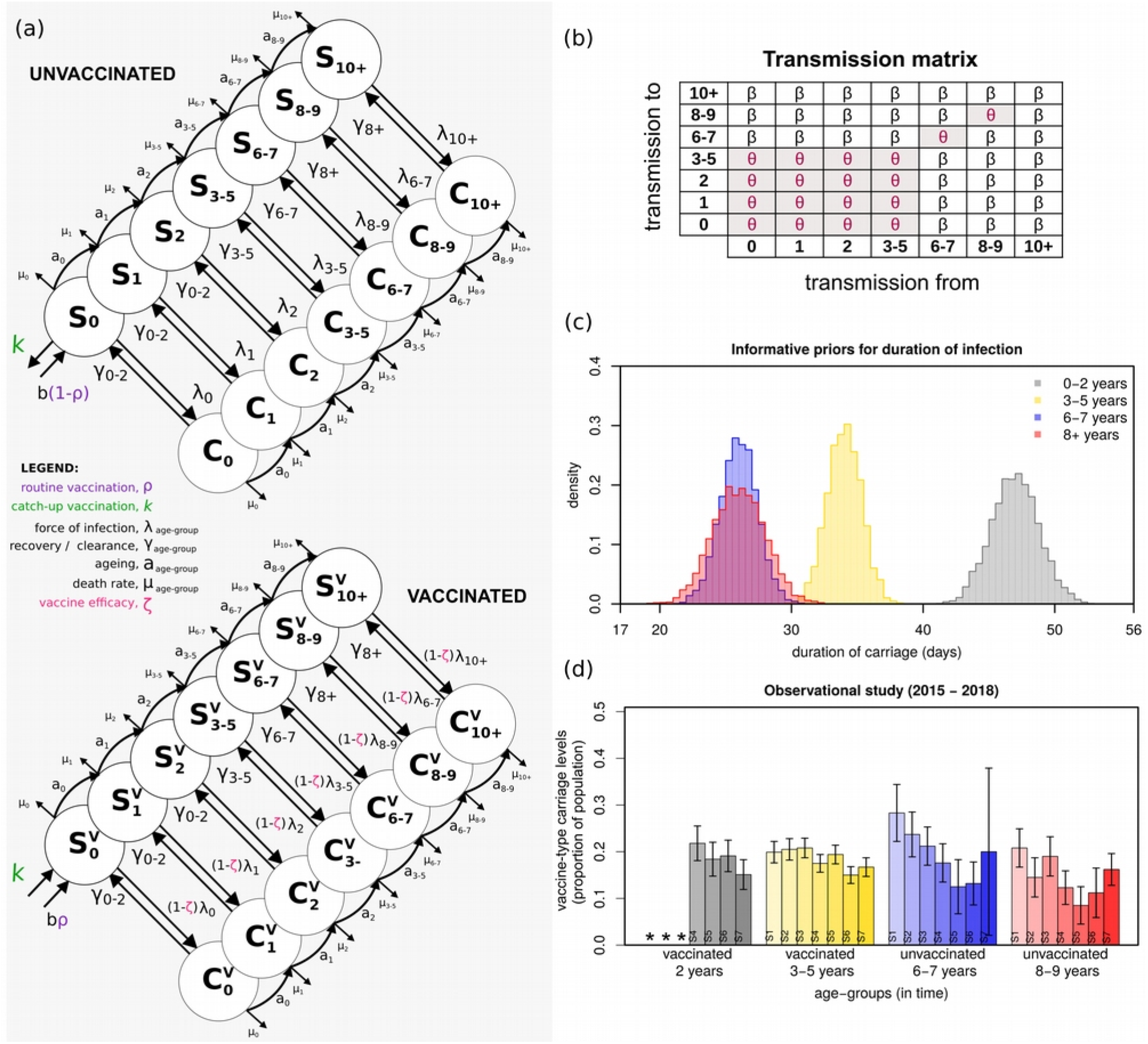
Survey data and model framework, priors and transmission matrix. **(a)** Seven age-groups were modelled: 0, 1, 2, 3-5, 6-7, 8-9, 10+ years of age (circles), each divided into unvaccinated (top) and vaccinated (bottom). Labels a_age-group_ mark ageing rates per age class; µ_age-group_ mark age-specific death rates; b marks births, at which point a proportion (ρ) are vaccinated) are vaccinated (purple); ζ marks vaccine-induced protection, expressed as reduction in susceptibility to infection of vaccinated individuals (magenta); λ_age-group_ mark age-specific forces of infection; γ_age-group_ mark age-specific rates of clearance from infection; k marks catch-up vaccination (green). **(b)** The transmission matrix used, with coefficients β and θ, where θ is the specific coefficient for transmission within and between particular age-groups. β and θ are estimated when fitting the survey data. **(c)** The informative priors used in the fitting exercise for mean (standard deviation) infectious periods (days) of 47 (1.8) for 0-2 years old; 34 (1.3) for 3-5 years old; 26 (1.4) for 6-8 years old; 26 (2.0) for 8+ years old (taken from [1]). The posterior values of these periods (1/γγ_0-2_, 1/γ γ_3-5_, 1/γγ_6-8_, 1/γγ_8+_) are estimated when fitting the survey data. **(d)** Mean and standard error for carriage as reported in the observational study data (surveys) per age-group (Table S7). S1 to S7 highlight the surveys 1 to 7. The * mark data that was not collected.

### Vaccine type transmission model

A deterministic, ordinary-differential equations (ODE) model (Figure 1a) was developed to fit VT carriage levels as reported in the cross-sectional observational study in Blantyre (Figure 1d)^22^. Fitting was implemented using a Bayesian Markov chain Monte Carlo (bMCMC) approach developed and used by us in other modelling studies^37–39^, including informative priors for duration of carriage (Figure 1b, Table S1) and uninformative uniform priors for vaccine efficacy (individual-level protection against carriage) and transmission potential. The methodology is summarised in this section and further details such as equations, literature review on priors and expected parameter values (Tables S1, S2, S5, S6) and complementary results can be found in Supplementary Text S1.

#### Pneumococcal infection dynamics and human demographics

As depicted in Figure 1a, the population was divided into seven non-overlapping age-groups: 0 (<1), 1, 2, 3-5, 6-7, 8-9, 10+ years old. Ageing was approximated by moving individuals along age-groups with a rate (a_age-group_) equal to the inverse of the time spent at each age class. The seven age-groups were further divided into vaccinated (S^v^_age-group_, C^v^_age-group_) and unvaccinated (S_age-group_, C_age-group_) susceptibles (S) and carriers (C). The population size was assumed to be constant, with total deaths equal to births (details in Supplementary Text S1). Death rates were age-specific (µ_age-group_) and relative to a generalized total life-span of 70 years.

#### Natural immunity

Pneumococcal colonization increases both humoral (anti-capsular serotype-specific and anti-protein non-serotype-specific) and T-cell (anti-protein) immunity^40^. Acquisition of this immunity correlates with colonization in children and increases with age as colonization decreases. In our model (Figure 1a), all individuals were assumed to be born susceptible but can acquire infection (colonization) at any age with a particular force of infection λ_age-group_, becoming carriers (C_age-group_) for an age-specific period (1/γ_age-group_), and returning to the susceptible state (S_age-group_) after clearance. Hence, the development of complete (sterile) immunity to the pneumococcus was not considered. We nonetheless allowed for decreasing duration of carriage with age (1/γ_age-group_) as a proxy for the development of pneumococcal immunity with age. To quantify differences in age, we used carriage duration data as reported by Hogberg and colleagues^41^ to define informative priors related to the aggregated age-groups: 0-2 years (1/γ_0-2_), 3-5 years (1/γ_3-5_), 6-8 years (1/γ_6-8_), and 8+ years (1/γ_8+_) as represented in Figure 1b (Table S1 for literature review).

#### Vaccination, efficacy and impact

For simplicity, routine vaccination was implemented at birth with coverage (*ρ*) at 92.5%^22^, and catch-up (*k*) implemented as a one-off transfer of a proportion of individuals from the unvaccinated susceptibles with 0 (<1) years of age (S_0_) to the vaccinated susceptible class with the same age (S^v^_0_) with coverage of 60%^22^. We assumed the vaccine to reduce the risk of infection (colonization) of vaccinated individuals by a proportion ζ (between 0 and 1, with ζ=1 equating to no risk). This reduction in risk was herein defined and interpreted as the individual-level vaccine efficacy against carriage (VE= 100 x ζ), and was modelled directly on the force of infection (λ) (Figure 1a, and Table S2 for literature review). We measured vaccine impact across age-groups as the post-PCV13 percent reduction in population-level VT carriage compared to pre-vaccination levels.

#### Force of infection

We considered several transmission matrices (Supplementary Text S1), and compared the resulting model fits using leave-one-out cross-validation (LOO) and the widely applicable information criterion (WAIC) measures. The inhomogeneous transmission matrix presented in Figure 1c over-performed the others and was used for the results presented in the main text. Its structure is based on epidemiological studies conducted in American, European and African populations reporting typical, strong, intrinsic variation in frequency, efficiency and environmental risk of transmission between age-groups^10,31,42–47^. In summary, the transmission matrix is generally populated with a baseline coefficient *β*, and a different coefficient *θ* assigned to transmission occurring within and between ages 0-5 years, and within 6-7 and 8-9 years of age independently. Further literature support and results from the second best performing transmission matrix can be found in Supplementary Text S1.

#### Fitting to survey data

The model’s carriage outputs for vaccinated 2, vaccinated 3-5, unvaccinated 6-7 and unvaccinated 8-9 years of age, were fitted to observed levels in Blantyre’s 1-7 surveys (Figure 1d, values in Table S7), approximately four to seven years PCV13 introduction (2015-2018). A total of seven parameters were fitted: vaccine efficacy against carriage (ζ, uninformative prior), coefficients of transmission (β, θ, uninformative priors) and durations of carriage in ages 0-2, 3-5, 6-7, 8+ years (1/γ_0-2_, 1/γ_3-5_, 1/γ_6-8_, 1/γ_8+_, informative priors). The transmission model was initialized at time t=0 with a proportion of 0.99 susceptibles and 0.01 infected, with numerical simulations run until an equilibrium was reached. At equilibrium, vaccination was introduced and the first post-vaccine 15 years recorded. Levels of carriage in the model were calculated as the proportion of individuals within an age-group that are carriers (i.e. *C/(S+C)*, expressions in Supplementary Text S1). The model was run with parameters scaled per year. bMCMC chains were run for 5 million steps, with burn-in of 20% (bMCMC details in see Supplementary Text S1).

## Results

We used our deterministic transmission model and bMCMC approach to fit the observed post-vaccination VT carriage data from Blantyre, Malawi (2015 - 2018). Based on this fit, we could reconstruct age-specific carriage dynamics for the unobserved first four years (2011 – 2015), and project VT carriage reduction into the future, to identify the mechanistic nature of the slow PCV13 impact on the vaccinated age-groups and strong herd-effects in the older unvaccinated age-groups.

### Model fit and posteriors

VT carriage levels across age-groups reported from the surveys were closely reproduced by the mean and 95% CI of the model using the bMCMC approach (Figure 2a). Our initial assumption of natural immunity accumulating with age was generally respected in the bMCMC solution (Figure 2b); i.e. the estimated posterior distributions of the durations of carriage (1/γ_age-group_) were adjusted by the bMCMC by approximately −0.7, +0.64, +0.58 and −1.73 days for the age-groups 0-2, 3-5, 6-7 and 8+ years of age, respectively. The posterior distribution of vaccine efficacy (individual-level protection against carriage) across ages was estimated to be 66.87% (95% CI 50.49-82.26). While we used an uninformative prior (uniform, 0 to 1) in the bMCMC, this efficacy posterior was similar to others recently estimated with different models and in multiple epidemiological settings (Figure 2c). We therefore argue that it serves as partial validation for our modelling framework. Finally, the solutions for the transmission coefficients *β* and *θ* suggested that in order to reproduce the Blantyre survey data, the risk of infection associated with contacts within and between younger age-groups (0-5 years old) would have to be higher than that of the general population (i.e. *θ*>>*β*).

**Figure 2:**
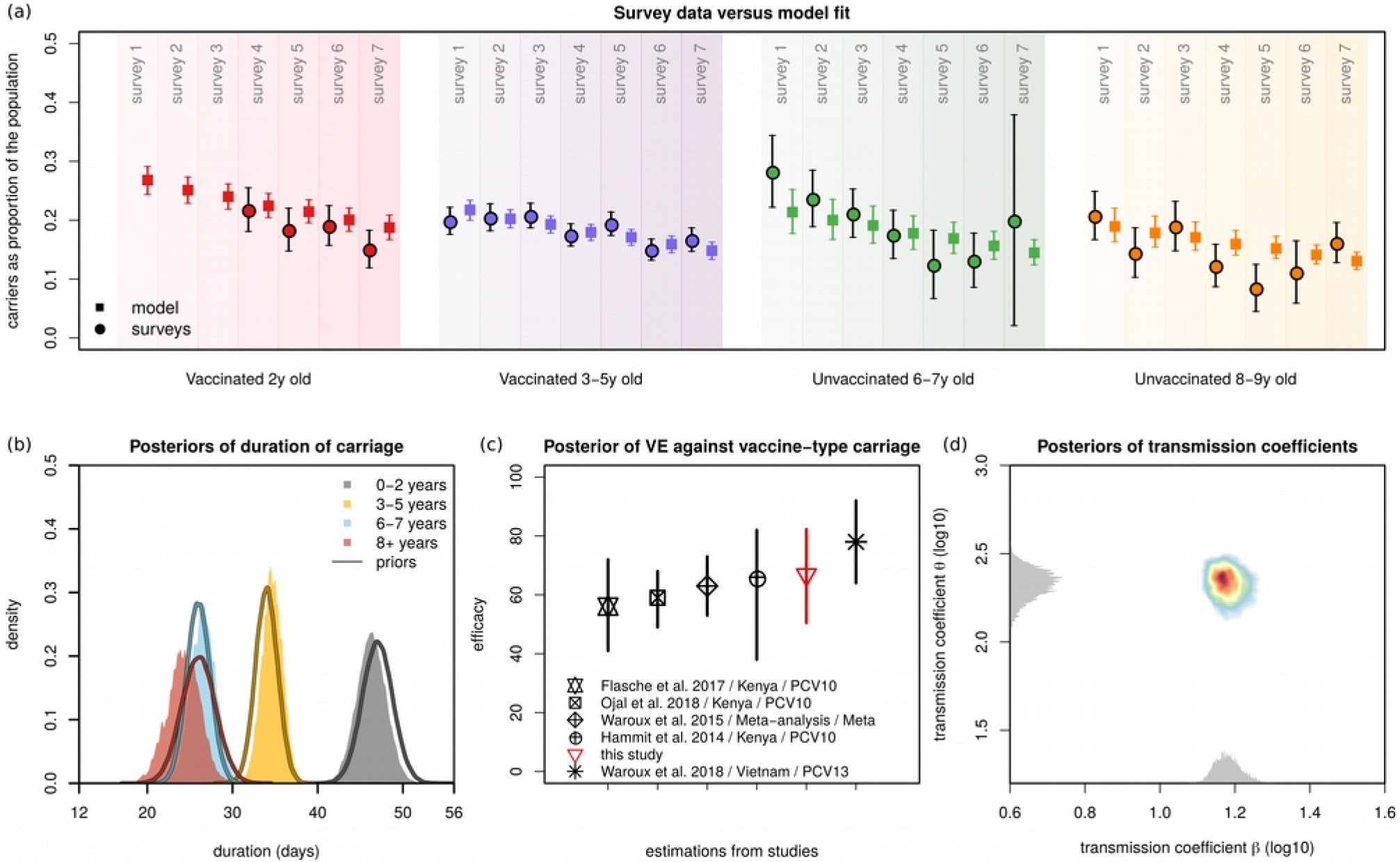
Model fit and estimated posteriors. **(a)** Model fit to carriage data from the observational study for different age-groups: vaccinated 2 years old (red), vaccinated 3-5 years old (purple), unvaccinated 6-7 years old (green) and unvaccinated 8-9 years old (orange). The survey data is represented by full circles, the model output by full squares (data in Figure 1d, Table S7). **(b)** Priors (lines) and estimated posterior distributions (shaded) of duration of carriage per age-group. **(c)** Estimated mean and 95% CI of posterior of vaccine efficacy against vaccine-type carriage (red) in the context of estimates from other studies (in legend, Table S2). **(d)** The estimated posterior distributions of the transmission coefficients β and θ are shown in two dimensions (coloured area). The estimated actual distribution for β is in the x-axis and θ in the y-axis (visualised in grey). Note that, for visualisation purposes, the axes are log_10_-transformed and the grey distributions’ height has no scale (height is not quantified). **(a,b,c,d)** Solutions presented are obtained from sampling 100,000 parameter values from posteriors and simulating the dynamic model.

### Vaccine impact across age-groups

Using parameter samples from the bMCMC estimated posteriors, we simulated vaccine impact in terms of VT carriage reduction across age-groups in the first 10 years post-vaccination (Figure 3).

**Figure 3:**
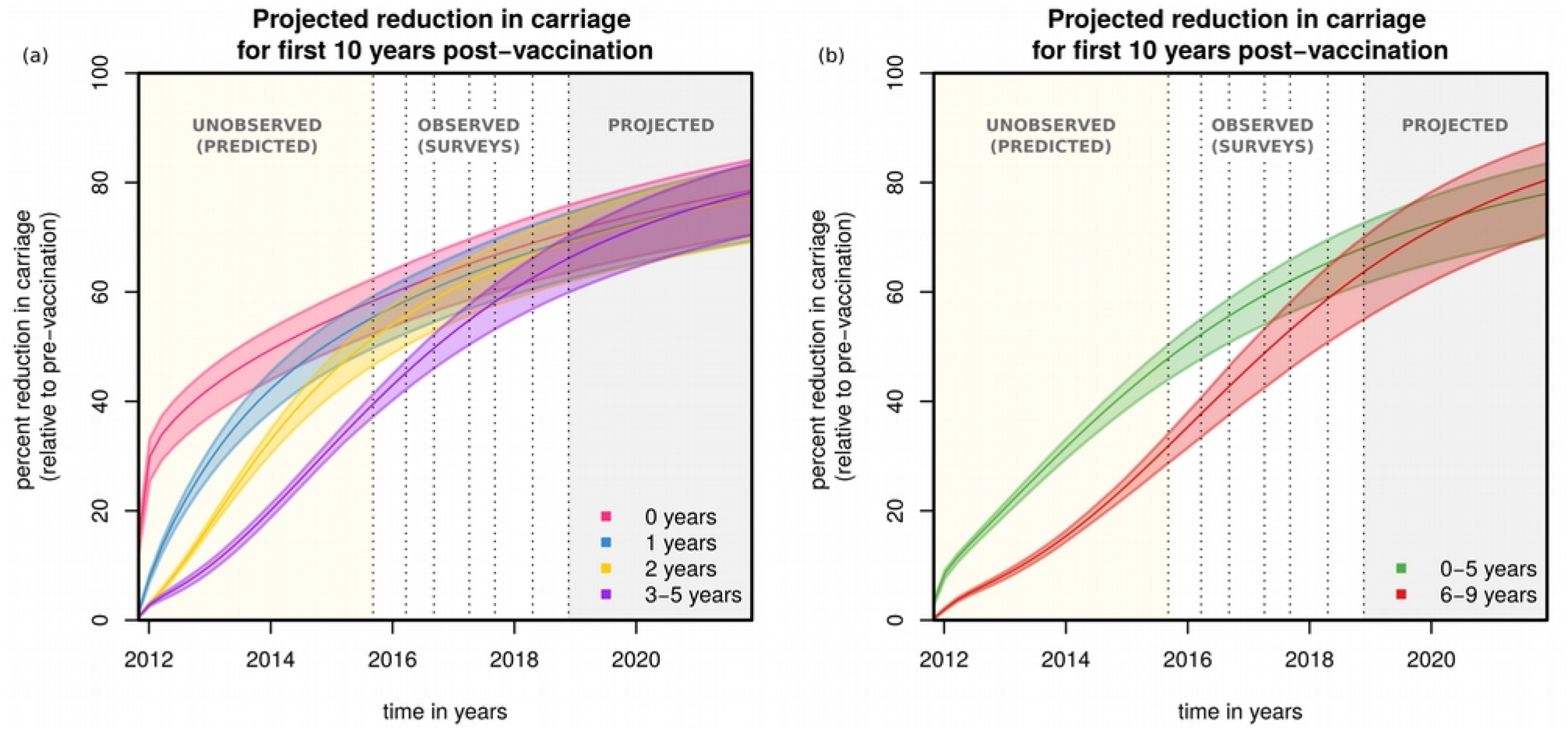
Projections of post-vaccination vaccine-type carriage reduction. **(a)** Projected reduction in carriage relative to the pre-vaccination era for age-groups 0 years (magenta), 1 year (blue), 2 years (yellow) and 3-5 years (purple) old. **(b)** Projected reduction in carriage relative to the pre-vaccination era for aggregated age-groups 0-5 years (green) and 6-9 years (red) old (with corresponding 95% CIs). **(a,b)** Solutions presented are obtained from sampling 100,000 parameter values from posteriors and simulating the dynamic model. The shaded areas are yellow for the post-vaccination period with no carriage data, white for the post-vaccination period with data, and grey for the post-vaccination projected period up to 10 years. Dotted vertical lines mark survey dates. The x-axis origin marks PCV13 introduction.

After the first year, VT carriage reduction was estimated to be 42.38% (95% CI 37.23-46.01%) for the 0 (<1) years old, followed by 29.25% (95% CI 26.4-31.4%) for the 1 years old, 17.45% (95% CI 16.47-18.36%) for the 2 years old and 4.95% (95% CI 8.78-10.89%) for 3-5 years old (Figure 3a). With time, as carriage generally dropped and vaccinated individuals aged, the older groups were estimated to benefit from increasingly similar reductions in carriage compared to the initially vaccinated group. Since during the first year only the 0 (<1) years of age were vaccinated, the short-term reductions in carriage of the other groups were due to indirect herd-effects alone.

At the target point of 10 years into the post-vaccination era, impact was estimated to be similar across all age-groups, with VT carriage reduced by 76.9% (CI 95% 68.93-82.32%) for the 0 (<1) years old, 75.72% (CI 95% 67.78-81.24%) for the 1 years old, 75.51% (CI 95% 67.55-81.05%) for the 2 years old and 75.86% (CI 95% 68.29-80.97%) for 3-5 years old. We further projected vaccine impact on aggregated age-groups 0-5 and 6-9 years of age, which showed equivalent reductions in VT carriage (Figure 3b), with the larger aggregated age-group 0-9 years old having a total reduction of 76.23% (CI 95% 68.02-81.96%) after 10 years.

We performed a literature review on observed reduction of VT carriage in time after the introduction of PCV vaccines (Table S5) in numerous countries, and concluded that both the observed carriage levels during the surveys and during the model’s projection for the first 10 years were high when compared to other countries. For instance, residual carriage of PCV13 types was 0.4% after 4 years of vaccination in England^48^, 9.1% after 2 years of vaccination in Italy^49^, and 7% after 3 years of vaccination in Alaska, USA^16^. Similarly, for 0-5 year old individuals, PCV10 in Kenya^18^ has reduced VT carriage by 73.92% in the first 5 years, while in Portugal^50^, PCV7 has reduced VT carriage by 78.91% in the same age-group and amount of time (more examples can be found on Table S5).

### Post-vaccination changes in force of infection

To try to understand responses to vaccination across age-groups, we further explored the post-PCV13 force of infection (FOI) dynamics. The FOI is the overall rate by which a certain age-group of susceptible individuals is infected, comprising the transmission rate (*β* or *θ*) weighted by the number of infectious individuals within the same and other age-groups. Although we modelled six independent age-groups under 10 years of age, only three unique FOIs are defined in the transmission matrix for individuals under 9 years of age (0-5, 6-7 and 8-9 years of age, Figure 1c).

As determined by the posteriors of *β* and *θ* (Figure 2d), the pre-vaccination absolute FOI of the 0-5, 6-7 and 8-9 age-groups was different at PCV13 introduction, and with vaccine roll out the FOI of each age-group decreased in time (Figure 4a). We also examined the FOI derivative with respect to time as a measure of speed of FOI reduction (Figure 4b), and found that the time period of fastest FOI reduction for the 0-5 years old was between vaccine introduction and 2015 (when no carriage data was collected). This contrasted with the older age-groups (6-7 and 8-9), for which the period of fastest FOI reduction was predicted to be just before or during the first three surveys. Thus, although surveys 1 to 7 suggest a rather slow reduction of VT carriage for the younger age-groups during the observational study, this seems to have been preceded by a period of high, short-term impact on VT carriage for those age-groups (seen in the initial dynamics of Figures 3a and 3b). Indeed, vaccine impact (reduction in VT carriage) at the time of the first survey was estimated to be 46.9% (95% CI 43.2-49.42) for the aggregated age-group 0-5 years old. At the same time, the fastest reduction in FOI for the older age-groups was predicted by the model to take place just before and during the first surveys, the time period in which survey data presents the largest reductions in VT carriage for those age-groups (Figure 1d). Overall, projected FOI dynamics suggest that PCV13 impact has been non-linear in time within age-groups, with predicted periods of faster reductions in VT carriage being experienced by different ages in a sequential manner, from younger to older individuals.

**Figure 4:**
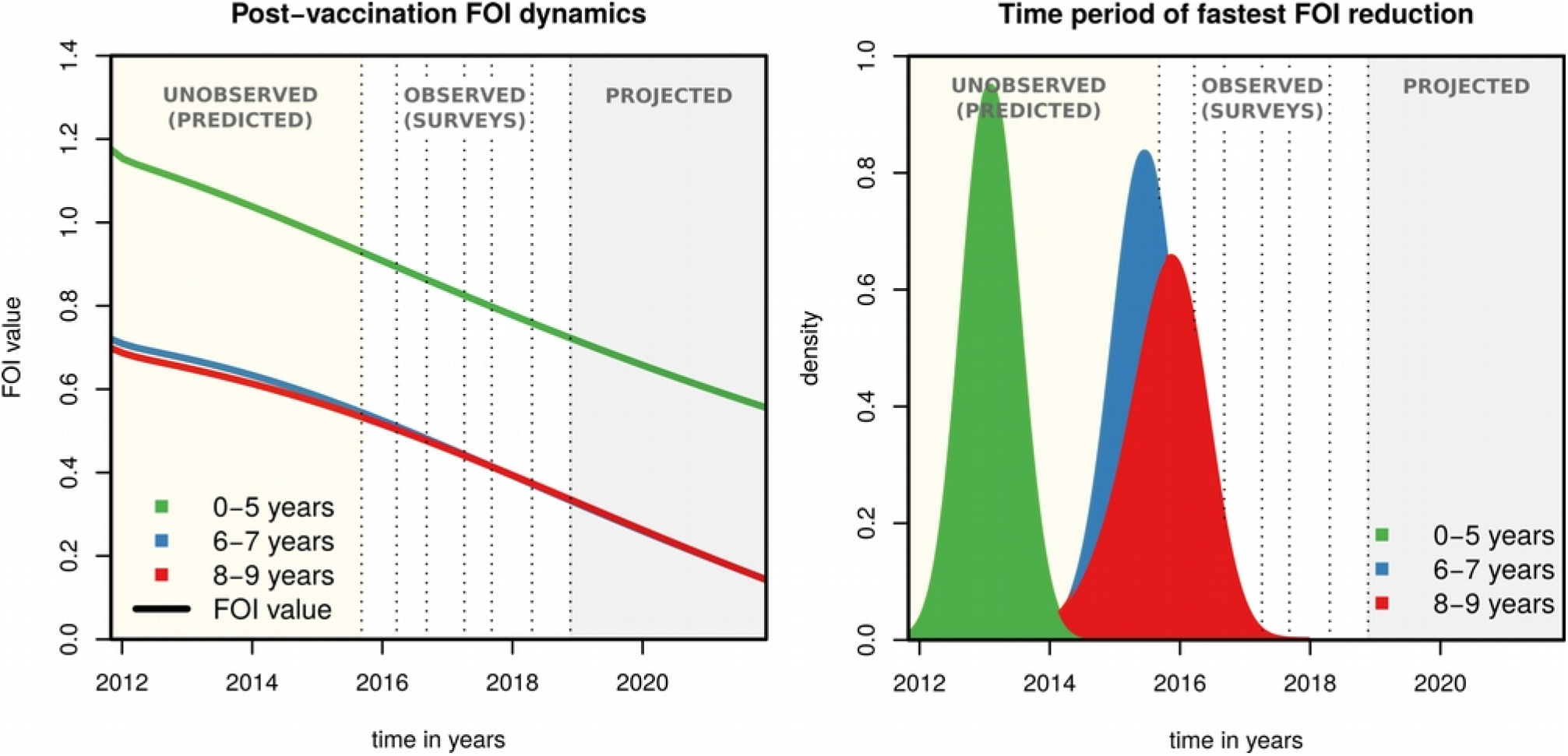
Projections of post-vaccination changes in the force of infection. **(a)** The post-vaccination force of infection (FOI) of different age groups (0-5 years in gree, 6-7 in blue and 8-9 in red) as calculated for each of 100,000 simulations using parameter samples from posteriors. **(b)** For each FOI of each age-group and each 100,000 simulations using parameter samples from posteriors, the time point of minimum derivative was calculated, resulting in one distribution per age-group (coloured curves, 0-5 years in green, 6-7 in blue, 8-9 in red). This time point is as a proxy for the period of fastest FOI reduction. The shaded areas are yellow for the post-vaccination period with no carriage data, white for the post-vaccination period with data, and grey for the post-vaccination projected period up to 10 years. Dotted vertical lines mark survey dates. The x-axis origin marks PCV13 introduction.

### Sensitivity of vaccine impact based on transmission setting

The projected impacts of Figures 3 and 4 were based on the estimated transmission coefficients for Blantyre (Figures 1b and 2d). To contextualize this particular transmission setting, we searched the literature for pre-vaccination VT carriage levels in other countries (Table S6). The reported age-groups were highly variable, and we therefore focused on the 0-5 years old group for which more data points were available from a range of countries in North America, Africa, Europe and South-east Asia (Figure 5a). Reported VT carriage in this age-group was highly variable both between and within countries, with our estimation for Blantyre being on the higher end (61.58%, 95% CI 50.0-70.9%).

**Figure 5:**
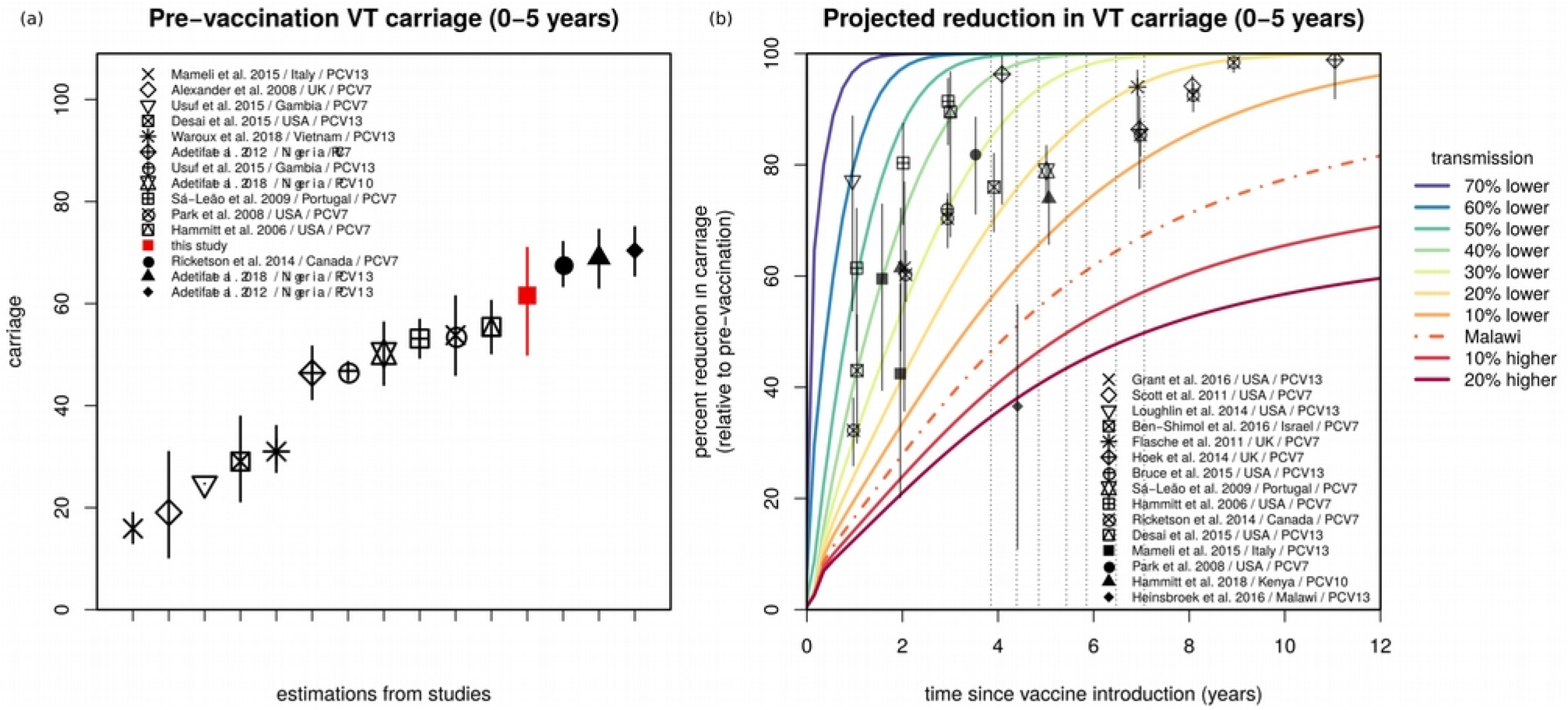
Estimated vaccine-type carriage and sensitivity of projections to baseline transmission in the context of other studies. **(a)** Estimated pre-vaccination vaccine-type carriage (and 95% CI) for the age-group 0-5 years of age (red) in the context of carriage levels reported in other studies (in legend, Table S6). **(b)** The baseline transmission coefficient (β) is varied by considering the 70%, 60%, 50%, 40%, 30%, 20%, and 10% lower, and 10%, 20% higher transmission than the estimated for Blantyre (Malawi, β_Malawi_) when fitting the observational study (e.g. 10% lower is 0.9*β_Malawi_). The impact projections for the age-group 0-5 years old using the β estimated for Blantyre (Malawi) are presented by the dashed line (as in Figure 3b). For visual purposes only the means are shown, obtained from simulations sampling 100,000 parameter values from posteriors. The symbols and whiskers are measures of reported impact (carriage reduction) and 95% CIs for several published studies (in legend, Table S5). The grey arrows mark the year of PCV13 introduction and the years of the four surveys.

We further searched the literature for post-vaccination VT carriage levels in other countries and again focused on the age-group 0-5 years old for which more data points were available (Table S5, points with whiskers in Figure 5b). The projected impact for Blantyre according to our model (dashed line), was notably lower than observed for other countries. A Malawi data point reported in the context of the Karonga District (Northern Malawi) had the closest impact to our projections in Blantyre (Southern Malawi), 4 to 5 years after PCV13 introduction^19^.

Given that our posterior of vaccine efficacy (individual-level protection against carriage, Figure 2c) was close to estimations from other regions of the world, we hypothesised that both the higher pre- and post-PCV13 VT carriage levels in Blantyre were likely due to a higher local force of infection compared to other regions. To demonstrate this, we simulated a range of alternative transmission settings in Blantyre, by varying both the transmission coefficients (*β* and *θ*) between −70% and +120% of their estimated posteriors (full exercise in Figure S3). This sensitivity exercise showed that lowering local transmission by approximately −30% was sufficient for the model to approximate short- and long-term vaccine impact observed in several other countries (Figure 5b). Other age-groups, for which far less data points were available, presented similar patterns (Figure S4).

## Discussion

Using a dynamic model, we have reproduced observed changes in pneumococcal VT carriage following the introduction of PCV13 in Blantyre, Malawi. Similar to other modelling frameworks we have considered the accumulation of natural immunity with age and have also allowed for heterogeneous transmission potentials within and between age-groups. Including these factors allowed us to identify age-related characteristics of the local force of infection as the main determinants of the high residual pneumococcal vaccine type carriage in Blantyre, seven years post-PCV13 introduction.

A main motivation for developing our dynamic model was to explain the high residual VT carriage levels seven years post-PCV13 introduction^22^. Studies from Kenya, The Gambia and South Africa have reported similar trends, with VT carriage remaining higher than in industrialised countries at similar post-vaccination time points. Compared to studies from other geographical regions, pre- and post-vaccination VT carriage in Blantyre was at the upper end of reported values across many countries (Figure 5 and Tables S5, S6). Given that our estimate of vaccine efficacy (individual-level protection against carriage) was similar to reports from elsewhere (Figure 2c, Table S2), we tested the hypothesis that the observed and projected lower vaccine impact was likely a result of a higher force of infection in Blantyre compared to other regions. This force of infection was found to be characterised by different transmission potentials within and between age-groups, and particularly dominated by individuals younger than 5 years. Reflecting a variety of approaches and assumptions that can be found in other models^8,11,28^, our framework is not able to discern if this assortative relationship with age is due to age-specific contact type patterns or susceptibility to colonization. Nonetheless, our results strongly argue for the need of more research characterising local contact, risk and transmission-route profiles (e.g. ^42^), if we are to understand the myriad of reported PCV impacts across different demographic, social and epidemiological settings.

There were also the observations of vaccine impact (reduction in VT carriage) in unvaccinated age-groups, and a particularly slow impact in younger vaccinated age-groups during the surveys (Figure 1d). The dynamic model helped explain these age-related responses, by showing that age-groups have experienced periods of higher vaccine impact at different time points, sequentially, from younger to older groups. A major implication is that reduction in VT carriage in vaccinated younger age-groups has been fastest between PCV13 introduction and 2015, when no carriage data was collected in Blantyre, but consistent with data collected in rural northern Malawi^19^. Thus, similarly to the conclusions of another modelling study^28^, our results advocate for the essential role of dynamic models to understand post-PCV13 VT carriage, by critically accounting for local non-linear effects of pneumococcal transmission and vaccination which may have significant implications for data interpretation.

Critical for low and middle income countries, as well as global initiatives such as Gavi, is that the impact of PCVs on pneumococcal VT carriage needs to be further improved if we are to maximize disease reduction. For high burden countries like Malawi, in which post-PCV VT carriage data suggests that local epidemiological factors may dictate lower vaccine impact than elsewhere, region-specific improved vaccination schedules^19,22^ and catch-up campaigns^28^ could help speed-up VT carriage reduction and maximise cost-effectiveness. For this to be possible, we need to better understand local transmission profiles across ages, which are likely dictated by demographic and socio-economic factors, and strongly determine short- and long-term PCV impact.

## Limitations

Data suggest that immune responses to PCV vaccines wane over time^22,34^. In a meta-analysis study, PCV7 efficacy was estimated at 62% (CI 95% 52-72%) at four months post-vaccination, decreasing to 57% (CI 95% 50-65%) at six months, but remaining 42% (CI 95% 19-54%) at five years post-vaccination^34^. Models implicitly parametrising for duration of vaccine-induced protection (dVP) have typically followed a prior with minimum mean duration of six years^8,11,28,34^, but in one study dVP was estimated as 8.3 years (95% CI 5 – 20)^8^. Our framework does not explicitly include dVP, and this should be a line of future modelling research. Due to the time ranges studied for Blantyre (data were collected up to seven years post-PCV13 introduction and projections made only up to the first ten years), we argue that our results should be robust and only weakly influenced by not considering dVP. In light of the possibility that dVP is shorter than previously reported^22^, our projections of vaccine impact should be seen as a best-case scenario; i.e. real long-term vaccine impact in Blantyre would likely be lower than projected by our model. Our framework also does not include niche competition between VT and non-VT pneumococci^11,28,34^. It is difficult to assert the impact of such competition in our main results, but it is unlikely that our conclusions would be significantly affected, since they are mostly based on factors which have not been reported to be associated with type competition directly (e.g. age-specific transmission).

## Conclusion

In Blantyre, vaccine efficacy (individual-level protection against carriage) across ages and time was estimated at 66.87% (95% CI 50.49-82.26%), similar to reports from other countries. However, local transmission potential in Blantyre is likely to be higher than in other countries and also heterogeneous among age-groups, with a particular contribution from younger children. While PCV13 is achieving positive outcomes in Blantyre^19,51^, a local higher and age-dependent force of infection is dictating a lower long-term vaccine impact (population-level carriage reduction) than reported elsewhere. Finally, the combination of age-related transmission heterogeneities and routinely vaccinating infants has led to non-linear responses in terms of vaccine impact across ages and time, with general implications on post-vaccination VT carriage data interpretation. Together, these findings suggest that in regions with lower than desired PCV impact on VT carriage, alternative vaccine schedules and catch-up campaigns targetting children <5 years of age should be further evaluated.

## Abbreviations

VT: vaccine type
NVT: non-vaccine type
PCV: pneumococcal conjugate vaccine
CI: confidence interval
bMCMC: Bayesian Markov chain Monte Carlo
ODE: ordinary-differential equations
FOI: force of infection
dVP: duration of vaccine-induced protection

## Acknowledgements

We would like to thank Ellen Heinsbroek for key literature references used in this manuscript, and Stefan Flasche for useful information regarding raw data of published studies. We thank the individuals who participated in this study and the local schools and authorities for their support. We are grateful to the study field teams (supported by Farouck Bonomali and Roseline Nyirenda) and the study laboratory team. We are grateful to the MLW laboratory management team (led by Brigitte Denis) and the MLW data management team (led by Clemens Masesa). RSH, NF and TS are supported by the National Institute for Health Research (NIHR) Global Health Research Unit on Mucosal Pathogens using UK aid from the UK Government. The views expressed in this publication are those of the author(s) and not necessarily those of the NIHR or the Department of Health and Social Care.

## Contributions

JL, UO, TDS designed the modelling study. JL and UO designed the model. JL implemented the model and the fitting approach. JL, UO analysed and interpreted model output. JL and UO searched and curated the literature data. TDS supervised, while AG, NBZ, DE, AWK, TSM, AAM, CM and MB collected and curated the Malawi observational data. SG, NF and RSH supervised both the modelling and observational sides of the study. JL wrote the first draft of the manuscript which all authors revised. JL, UO and TDS revised other iterations of the manuscript. All authors revised the last version of the manuscript.

