## Supplementary Material for "Determinants of high residual post-PCV13 pneumococcal vaccine type carriage in Blantyre, Malawi: a modelling study"

Supplementary Text S1: methodological details, literature support and complimentary results.

### Ordinary-differential equations (ODE)

Equations for the vaccinated (equations 1-14) and unvaccinated (equations 15-28) age-groups, with model diagram in Figure 1 of main text:

$$
\begin{aligned}
dS_0/d_t &= b(1-\rho) - \lambda_0 S_0 + \gamma_{0-2} C_0 - a_0 S_0 - \mu_0 S_0 && (1)\\
dC_0/d_t &= \lambda_0 S_0 - \gamma_{0-2} C_0 - a_0 C_0 - \mu_0 C_0 && (2)\\
dS_1/d_t &= a_0 S_0 - \lambda_1 S_1 + \gamma_{0-2} C_1 - a_1 S_1 - \mu_1 S_1 && (3)\\
dC_1/d_t &= a_0 C_0 + \lambda_1 S_1 - \gamma_{0-2} C_1 - a_1 C_1 - \mu_1 C_1 && (4)\\
dS_2/d_t &= a_1 S_1 - \lambda_2 S_2 + \gamma_{0-2} C_2 - a_2 S_2 - \mu_2 S_2 && (5)\\
dC_2/d_t &= a_1 C_1 + \lambda_2 S_2 - \gamma_{0-2} C_2 - a_2 C_2 - \mu_2 C_2 && (6)\\
dS_{3-5}/d_t &= a_2 S_2 - \lambda_{3-5} S_{3-5} + \gamma_{3-5} C_{3-5} - a_{3-5} S_{3-5} - \mu_{3-5} S_{3-5} && (7)\\
dC_{3-5}/d_t &= a_2 C_2 + \lambda_{3-5} S_{3-5} - \gamma_{3-5} C_{3-5} - a_{3-5} C_{3-5} - \mu_{3-5} C_{3-5} && (8)\\
dS_{6-7}/d_t &= a_{3-5} S_{3-5} - \lambda_{6-7} S_{6-7} + \gamma_{6-7} C_{6-7} - a_{6-7} S_{6-7} - \mu_{6-7} S_{6-7} && (9)\\
dC_{6-7}/d_t &= a_{3-5} C_{3-5} + \lambda_{6-7} S_{6-7} - \gamma_{6-7} C_{6-7} - a_{6-7} C_{6-7} - \mu_{6-7} C_{6-7} && (10)\\
dS_{8-9}/d_t &= a_{6-7} S_{6-7} - \lambda_{8-9} S_{8-9} + \gamma_{8+} C_{8-9} - a_{8-9} S_{8-9} - \mu_{8-9} S_{8-9} && (11)\\
dC_{8-9}/d_t &= a_{6-7} C_{6-7} + \lambda_{8-9} S_{8-9} - \gamma_{8+} C_{8-9} - a_{8-9} C_{8-9} - \mu_{8-9} C_{8-9} && (12)\\
dS_{10+}/d_t &= a_{8-9} S_{8-9} - \lambda_{10+} S_{10+} + \gamma_{8+} C_{10+} - \mu_{10+} S_{10+} && (13)\\
dC_{10+}/d_t &= a_{8-9} C_{8-9} + \lambda_{10+} S_{10+} - \gamma_{8+} C_{10+} - \mu_{10+} C_{10+} && (14)
\end{aligned}
$$

$$
\begin{aligned}
dS^v_0/d_t &= b\rho - (1-\zeta)\lambda_0 S^v_0 + \gamma_{0-2}C^v_0 - a_0 S^v_0 - \mu_0 S^v_0 && (15)\\
dC^v_0/d_t &= (1-\zeta)\lambda_0 S^v_0 - \gamma_{0-2}C^v_0 - a_0 C^v_0 - \mu_0 C^v_0 && (16)\\
dS^v_1/d_t &= a_0 S^v_0 - (1-\zeta)\lambda_1 S^v_1 + \gamma_{0-2}C^v_1 - a_1 S^v_1 - \mu_1 S^v_1 && (17)\\
dC^v_1/d_t &= a_0 C^v_0 + (1-\zeta)\lambda_1 S^v_1 - \gamma_{0-2}C^v_1 - a_1 C^v_1 - \mu_1 C^v_1 && (18)\\
dS^v_2/d_t &= a_1 S^v_1 - (1-\zeta)\lambda_2 S^v_2 + \gamma_{0-2}C^v_2 - a_2 S^v_2 - \mu_2 S^v_2 && (19)\\
dC^v_2/d_t &= a_1 C^v_1 + (1-\zeta)\lambda_2 S^v_2 - \gamma_{0-2}C^v_2 - a_2 C^v_2 - \mu_2 C^v_2 && (20)\\
dS^v_{3-5}/d_t &= a_2 S^v_2 - (1-\zeta)\lambda_{3-5} S^v_{3-5} + \gamma_{3-5}C^v_{3-5} - a_{3-5} S^v_{3-5} - \mu_{3-5} S^v_{3-5} && (21)\\
dC^v_{3-5}/d_t &= a_2 C^v_2 + (1-\zeta)\lambda_{3-5} S^v_{3-5} - \gamma_{3-5}C^v_{3-5} - a_{3-5} C^v_{3-5} - \mu_{3-5} C^v_{3-5} && (22)\\
dS^v_{6-7}/d_t &= a_{3-5} S^v_{3-5} - (1-\zeta)\lambda_{6-7} S^v_{6-7} + \gamma_{6-7}C^v_{6-7} - a_{6-7} S^v_{6-7} - \mu_{6-7} S^v_{6-7} && (23)\\
dC^v_{6-7}/d_t &= a_{3-5} C^v_{3-5} + (1-\zeta)\lambda_{6-7} S^v_{6-7} - \gamma_{6-7}C^v_{6-7} - a_{6-7} C^v_{6-7} - \mu_{6-7} C^v_{6-7} && (24)\\
dS^v_{8-9}/d_t &= a_{6-7} S^v_{6-7} - (1-\zeta)\lambda_{8-9} S^v_{8-9} + \gamma_{8+}C^v_{8-9} - a_{8-9} S^v_{8-9} - \mu_{8-9} S^v_{8-9} && (25)\\
dC^v_{8-9}/d_t &= a_{6-7} C^v_{6-7} + (1-\zeta)\lambda_{8-9} S^v_{8-9} - \gamma_{8+}C^v_{8-9} - a_{8-9} C^v_{8-9} - \mu_{8-9} C^v_{8-9} && (26)\\
dS^v_{10+}/d_t &= a_{8-9} S^v_{8-9} - (1-\zeta)\lambda_{10+} S^v_{10+} + \gamma_{8+}C^v_{10+} - \mu_{10+} S^v_{10+} && (27)\\
dC^v_{10+}/d_t &= a_{8-9} C^v_{8-9} + (1-\zeta)\lambda_{10+} S^v_{10+} - \gamma_{8+}C^v_{10+} - \mu_{10+} C^v_{10+} && (28)
\end{aligned}
$$

### Definition of birth rate for a constant population size

Given that the total population size is kept constant, the birth rate is a composite expression of deaths across ages:

$$
\begin{aligned}
b = {} & \mu_0(S_0 + C_0 + S^v_0 + C^v_0) + \\
& \mu_1(S1 + C1 + S^v_1 + C^v_1) + \\
& \mu_2(S2 + C2 + S^v_2 + C^v_2) + \\
& \mu_{3-5}(S_{3-5} + C_{3-5} + S^v_{3-5} + C^v_{3-5}) + \\
& \mu_{6-7}(S_{6-7} + C_{6-7} + S^v_{6-7} + C^v_{6-7}) + \\
& \mu_{8-9}(S_{8-9} + C_{8-9} + S^v_{8-9} + C^v_{8-9}) + \\
& \mu_{10+}(S_{10+} + C_{10+} + S^v_{10+} + C^v_{10+}) \qquad (29)
\end{aligned}
$$

### Expressions for model carriage levels

Levels of carriage in the ODE model are calculated as the proportion of individuals within an age-group, with a particular vaccine status, that are carriers at the time of the surveys. For example:

$$
\begin{aligned}
Ca_0 &= C_0 + C^v_0 && (37)\\
Ca_1 &= C_1 + C^v_1 && (38)\\
Ca_2 &= C_2 + C^v_2 && (39)\\
Ca_{3-5} &= C_{3-5} + C^v_{3-5} && (40)\\
Ca_{6-7} &= C_{6-7} + C^v_{6-7} && (41)\\
Ca_{8-9} &= C_{8-9} + C^v_{8-9} && (42)\\
Ca_{10+} &= C_{10+} + C^v_{10+} && (43)\\
& && (44)
\end{aligned}
$$

### Markov-chain Monte-Carlo fitting approach

We used a Bayesian Markov chain Monte Carlo (bMCMC) approach, developed and used by us in other modelling studies [1]–[3]. The proposal distributions (*q*) of each parameter are defined as Gaussian (symmetric), effectively implementing a random walk Metropolis kernel. We define our acceptance probability $\alpha$ of a parameter set $\Theta$ given model ODE output y as:

$$\alpha = min\{1, \frac{\pi(y|\Theta^\star)p(\Theta^\star)q(\Theta^o|\Theta^\star)}{\pi(y|\Theta^o)p(\Theta^o)q(\Theta^\star|\Theta^o)}\} \qquad (45)$$

where $\Theta^*$ and $\Theta^o$ are the proposed and current (accepted) parameter sets (respectively); $\Pi(y \mid \Theta^*)$ and $\Pi(y \mid \Theta^o)$ are the likelihoods of the ODE output representing the (fitted) variables by each parameter set; $p(\Theta^o)$ and $p(\Theta^*)$ are the prior-related probabilities given each parameter set.

For simplicity and because all fitted variables are proportions, the likelihoods π were calculated as the product of conditional Gaussian probabilities (*Pr{...}*). The likelihood is the product the conditional probabilities of all variables, and can be formally expressed as:

$$\pi(y|\Theta) = \prod_{i=1}^{N} [Pr\{y_i = d_i\}] \qquad (46)$$

### Literature support for estimated durations of carriage

While we base our priors on the study by Hogberg and colleagues [4], there is vast support in the literature for both the use of similar values and the key assumption of a decrease in duration of carriage with age. We summarize this support in Table S1 below.

| Age-group | Duration | Country | Ref. |
|---|---|---|---|
| 0-1 years | 71 | UK | [10]–[12] |
| 2-4 years | 28 | UK | [10]–[12] |
| 5-17 years | 18 | UK | [10]–[12] |
| 18+ years | 17 | UK | [10]–[12] |
| 0-1 years | 30 | Sweden | [13] |
| 1-4 years | 21 | Sweden | [13] |
| 5-6 years | 13 | Sweden | [13] |
| 7-18 years | 15 | Sweden | [5] |
| 18+ years | 14 | Sweden | [5] |
| 0-2 years | 60* | Thailand & Myanmar | [6] |
| Mothers | 31* | Thailand & Myanmar | [6] |
| 6-11 months | 48.5* | Kenya | [7] |
| 3-59 months | 31.3* | Kenya | [7] |
| 0-7 years | 44* | Finland | [8] |
| 7+ years | 36 | Finland | [8] |

| | | | |
|---|---|---|---|
| 0-5 years | 42.6* | Bangladesh | [9] |
| 5+ years | 38 | Bangladesh | [9] |

**Table S1 – Support for decrease of duration of carriage with age.** Mean values are presented and * marks values for which 95% CI are available in the original study. Grey and white backgrounds are a visual cue for different countries.

### Literature support for estimated vaccine efficacy against carriage

The posterior for vaccine efficacy against carriage (ζ) was shown in the main text to be in the range of estimates obtained from other studies. Table S2 presents the values reported in such studies.

| VE | Vaccine | Ref. | Country / Region |
|---|---|---|---|
| 56% (95% CI 41 - 72) | PCV10 | [14] | Kenya |
| 66% (95% CI 38 - 82) | PCV10 | [15] | Kenya |
| 63% (95% CI 53 - 73) | Meta | [16] | Meta-analysis |
| 78% (95% CI 64 - 92) | PCV13 | [17] | Vietnam |
| 59% (95% CI 49 - 68) | PCV10 | [18] | Kenya |

**Table S2 – Support for estimated vaccine efficacy.** Mean values, 95% CI, vaccine, reference and country are presented for each of the studies. These quantifications were reported in the original articles (were not estimated by us on reported data).

### Literature support and model sensitivity to different transmission matrices

We formulated five different transmission matrices (Figure S1). One matrix assumed that transmission potential between age-groups is homogeneous (Figure S1e), while the other four assumed variations of inhomogeneous transmission potential between the groups (Figure S1a-d).

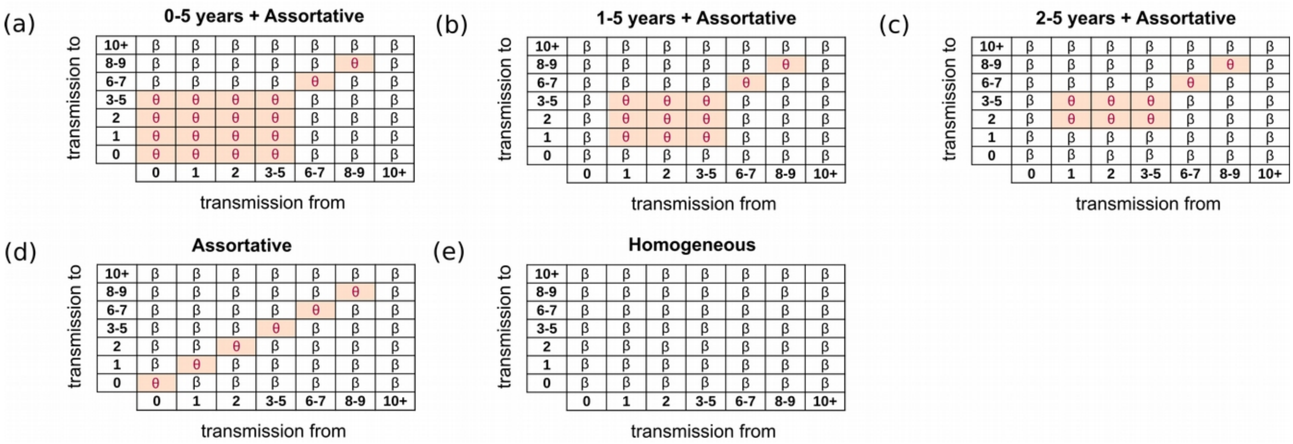

**Figure S1: Transmission matrices used for sensitivity of the model.** Subplots (a) to (e) present the five transmission matrices used in five independent fitting exercises of the model to the survey data described in the main text. The names of each matrix (and model) is above each subplot. The β and θ letters refer to the transmission coefficients described in the main text. The coefficient θ is in principle larger than β, but this relationship is not explicitly forced in the fitting approach.

The assumption, observation or estimation that pneumococcal transmission is inhomogeneous across ages is common in the literature. For instance, Ojal *et al.* have estimated that the probabilities of acquisition (colonization) across ages in Kilifi, Kenya after the introduction of PCV10, concluding that the probability of the age-group 1-5 years old is about 1.25 and 5.7 times higher than in the age-groups 6-14 and 15-20 years old [19]. For Finland [20], it has been estimated that around 44% of colonization events have a child <7 years of age as their source and that 18% of transmission events occured within this age group; and in Italy, the odds ratio for colonization in children aged <2 years after PCV13 introduction was still high at 3.75 [21]. In a different study by Adetifa *et al.*, carriage was 1.4 times higher in children <4 years of age compared to the next age-group 5-14 years of age [22]. Still in Africa, Uganda, Polain de Waroux *et al.* found that the age-group 5-14 years had the highest frequency of contacts, and a strong age-assortative pattern was seen across age-groups [23]. The latter effect has been reported in several countries including Uganda, Kenya, Finland, USA, and Great Britain [20], [23]–[26]. The epidemiological importance of considering higher efficiency and / or contact numbers between and within younger age-groups is not restricted to the pneumococcus, and is often reported for other pathogens as well [27].

After fitting the model to the survey data in five independent exercises (each with one of the matrices in Figure S1), we compared the frameworks using leave-one-out cross-validation (LOO) and widely applicable information criterion (WAIC) measures. Comparisons were done based on (1) the estimated posteriors versus priors, and (2) posterior model carriage levels versus observed surveys 1 – 7. The priors included were the carriage duration by Hogberg and colleagues [4] (Table S1, Figure 1b) and the vaccine efficacy against carriage (Table S2 and Figure 2c). LOO and WAIC are methods that allow estimating pointwise out-of-sample prediction accuracy from the five models using the log-likelihood evaluated at simulations from the parameter values of the estimated posteriors [28]. We used the loo R-package [29] to calculate model weights: 1) WAIC weights, 2) Pseudo-BMA weights without Bayesian bootstrap, 3) Pseudo-BMA + weights with Bayesian bootstrap, and 4) Bayesian stacking weights [30]. All of these measures vary between 0 and 1, with the sum between models adding up to 1; the highest the weight, the more favoured a model is compared to the rest. That is, a higher weight points to a more balanced response from the model, both in terms of reproducing the survey data but also in respecting the priors for duration of carriage and the literature knowledge on vaccine efficacy against carriage. The results of model comparison are in Table S3.

| **model / matrix** | **1) WAIC** | **2) Pseudo-BMA** | **3) Pseudo-BMA with BS** | **4) Bayesian stacking** |
|---|---|---|---|---|
| 0-5 years +Assortative | 0.69 | 0.99 | 0.87 | 0.84 |
| 1-5 years + Assortative | 0.31 | 0.01 | 0.13 | 0.16 |
| 2-5 years + Assortative | 0.00 | 0.00 | 0.00 | 0.00 |
| Assortative | 0.00 | 0.00 | 0.00 | 0.00 |
| Homogeneous | 0.00 | 0.00 | 0.00 | 0.00 |

**Table S3 – LOO and WAIC weights between model frameworks.**

The best score was for the model with the 0-5 years + Assortative transmission matrix (Figure 1a). This is therefore the model presented in the main text. In this model, the transmission potential (θ) among individuals 0-5 years old, and among 7-6 and 8-9 years old is different (larger, Figure 2d) than among other age-groups (β). The second best scored model (1-5 years + Assortative, Figure 1b) was the one using the most similar matrix to 0-5 years + Assortative, but which critically did not consider individuals of age 0 years old (i.e. <1 years). For completion, we present the general output of the 1-5 years + Assortative model in Figure S2.

| age-group | Carriage reduction after 1 year |
|---|---|
| **0** | 48.2 (38.7-57.1) |
| **1** | 26.87 (23.1-29.7) |
| **2** | 14.9 (13.4-15.9) |
| **3-5** | 7.3 (6.6-7.8) |
| **0-5** | 16.5 (14.9-17.1) |
| **6-9** | 6.1 (5.6-6.7) |

| Vaccine efficacy |
|---|
| 62.28 (46.24-77.60) |

| age-group | Carriage reduction after 10 years |
|---|---|
| **0** | 79.8 (67.9-87.7) |
| **1** | 75.1 (65.2-81.7) |
| **2** | 74.3 (64.4-80.7) |
| **3-5** | 74.6 (64.9-80.8) |
| **0-9** | 75.1 (64.5-81.9) |

| Pre-vaccination carriage 0-5 years |
|---|
| 51.2 (41.3-59.3) |

| age-group | Posterior adjustment to priors of carriage duration |
|---|---|
| **0-2** | -0.68 |
| **3-5** | +0.58 |
| **6-7** | +0.53 |
| **8+** | -1.71 |

| age-group | 10 year FOI reduction |
|---|---|
| **0-5** | 5.08 |
| **6-7** | 6.09 |
| **8-9** | 5.93 |

**Table S4 – Posteriors for the model 1-5 years + Assortative.** Ranges are the 95% CI.

The 1-5 years + Assortative model presents differences in output and capacity to fit the data. For instance, large differences in the posterior for vaccine efficacy against carriage and the estimation of pre-vacination VT carriage among 0-5 years old can be visually identified comparing Figure 2c, Figure 5a and Figure S2. For completion, we include here the 1-5 years + Assortative model full range of results (Table S4) presented in the main text for the 0-5 years + Assortative.

Although we defined the model transmission matrices similarly to matrices reported in previous studies, it is likely that further important heterogeneities exist between age-groups. To assess this, however, would require the addition of several parameters to the system, which in turn would increase the complexity and uncertainty of the fitting exercise. It is also the case that without further empirical support, it is difficult to plan and develop this new parametrization, as the required details on which age-group have specific lower / higher transmission coefficients in the particular context of the Blantyre cohort is largely unknown.

The independent contribution of either mixing (e.g. contact frequency) or risk (e.g. efficiency of contact) to the force of infection of each age-group can not be disentangled in our modelling framework. Nonetheless, we note that both higher mixing and risk of infection are supported in our study. For the first, as described above, we resort to literature reports on particular mixing patterns between age groups. For the second, we note that a higher proportion of children 3-6 years of age lived in houses with some lower quality infrastructure and had greater reliance on shared communal water sources in the cohort of our observational study [31].

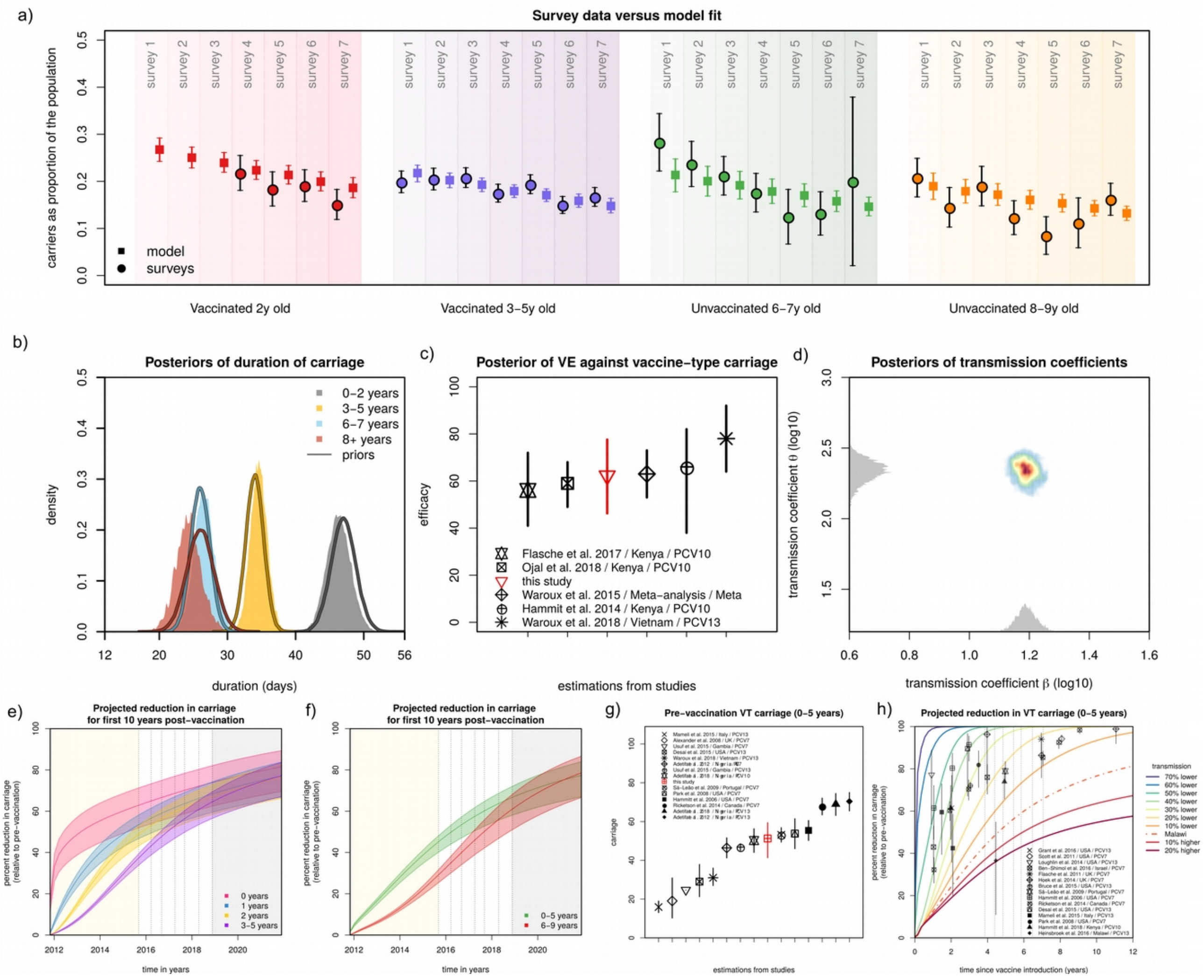

**Figure S2: Model fit and estimated posteriors when using a framework that uses the 1-5 years + Assortative transmission matrix. (a)** Model fit to carriage data from the observational study for age-groups: vaccinated 2 years old (red), vaccinated 3-5 years old (purple), unvaccinated 6-7 years old (green) and unvaccinated 8-9 years old (orange). The survey data is represented with means as full circles, the model output with means as full squares; the whiskers are the 95% CI. **(b)** Priors (lines) and estimated posterior distributions (shaded areas) for duration of carriage per age-group. **(c)** Estimated posterior for vaccine efficacy (and 95% CI) against vaccine-type carriage (red) in the context of estimations from other studies (in legend, Table S2). **(d)** Estimated posterior for the transmission coefficient β. **(e)** Projected reduction in carriage relative to the pre-vaccination era for age-groups 0 years (magenta), 1 year (blue), 2 years (yellow) and 3-5 years (purple) old. **(f)** Projected reduction in carriage relative to the pre-vaccination era for aggregated age-groups 0-5 years (green) and 6-9 years (red) old (with corresponding 95% CIs). The shaded areas are: yellow for the post-vaccination period with no carriage data, white for the post-vaccination period with survey carriage data, and grey for the post-vaccination projected period up to 10 years. **(g)** Estimated pre-vaccination vaccine-type carriage (and 95% CI) for the age-group 0-5 years of age (red) in the context of carriage levels reported in other studies (in legend, Table S6). **(h)** The baseline transmission coefficient (β) is varied by considering the 70%, 60%, 50%, 40%, 30%, 20%, and 10% lower, and 10%, 20% higher transmission than the estimated for Blantyre (Malawi, $\beta_{Malawi}$) when fitting the observational study (e.g. 10% lower is $0.9*\beta_{Malawi}$). The impact projections for the age-group 0-5 years old using the β estimated for Blantyre (Malawi) are presented by the dashed line (as in Figure 3b). The symbols and whiskers are measures of reported impact (carriage reduction) and 95% CIs for several published studies (in legend, Table S5). **(a,b,c,d,e,f,g,h)** Solutions presented are obtained from sampling 100,000 parameter values from posteriors and simulating the dynamic model.

### Literature review of carriage reduction in time

The pre- and post-vaccination proportions of vaccine-type carriage were used to calculate a Relative Risk (RR) and its corresponding 95% CI (under asymptotic normality). Using the expression (carriage(post) – carriage(pre)) / carriage(post) = 1 – carriage(pre) / carriage(post) = 1 - RR yielded a corresponding 95% CI for the relative change in carriage (Table S5) for particular time points after vaccination, age-groups, vaccines and countries. These values are used in figures of the main text and other supplementary figures comparing results from Blantyre (Malawi) with other countries.

| Percent Reduction | Time | Age | Vaccine | Ref. | Country / Region |
|---|---|---|---|---|---|
| 80.13% (95% CI 70.31 - 86.71) | 3 | 11 m | PCV7 | [36] | Netherlands |
| 69.33% (95% CI 46.65 - 82.37) | 4 | 6–23 m | PCV7 | [43] | USA |
| 34.84% (95% CI 24.52 - 43.74) | 2 | 6–24 m | PCV7 | [50] | France |
| 52.62% (95% CI 43.6 - 60.2) | 3 | 6–24 m | PCV7 | [50] | France |
| 60.88% (95% CI 53.27 - 67.26) | 4 | 6–24 m | PCV7 | [50] | France |
| 49.63% (95% CI 33.99 - 61.57) | 1 | 6–24 m | PCV13 | [51] | France |
| 95.31% (95% CI 93.44 - 96.65) | 5 | <2 y | PCV13 | [52] | China |
| 88.05% (95% CI 79.64 - 92.99) | 3 | 2 y | PCV7 | [36] | Netherlands |
| 77.17% (95% CI 53.76 - 88.73) | 1 | <5 y | PCV13 | [32] | USA |
| 78.91% (95% CI 73.16 - 83.42) | 5 | <5 y | PCV7 | [33] | Portugal |
| 61.44% (95% CI 46.63 - 72.14) | 1 | <5 y | PCV7 | [34] | USA |
| 80.32% (95% CI 69.28 - 87.39) | 2 | <5 y | PCV7 | [34] | USA |
| 91.41% (95% CI 83.65 - 95.49) | 3 | <5 y | PCV7 | [34] | USA |
| 32.24% (95% CI 25.93 - 38.02) | 1 | <5 y | PCV7 | [35] | Canada |
| 60.26% (95% CI 55.48 - 64.52) | 2 | <5 y | PCV7 | [35] | Canada |
| 70.36% (95% CI 65.14 - 74.79) | 3 | <5 y | PCV7 | [35] | Canada |
| 85.35% (95% CI 81.62 - 88.32) | 7 | <5 y | PCV7 | [35] | Canada |
| 92.52% (95% CI 89.6 - 94.61) | 8 | <5 y | PCV7 | [35] | Canada |
| 98.41% (95% CI 96.67 - 99.24) | 9 | <5 y | PCV7 | [35] | Canada |
| 89.56% (95% CI 67.11 - 96.69) | 3 | <5 y | PCV13 | [37] | USA |
| 42.45% (95% CI 20.15 - 58.52) | 2 | <5 y | PCV13 | [38] | Italy |
| 59.5% (95% CI 39.48 - 72.9) | 1.5 | <5 y | PCV13 | [38] | Italy |
| 81.82% (95% CI 71.19 - 88.52) | 3.5 | <5 y | PCV7 | [39] | USA |
| 94% (95% CI 84 - 97) | 7 | <5 y | PCV7 | [40] | UK |
| 43% (95% CI 30 - 53) | 1 | <5 y | PCV7 | [41] | Israel |
| 76% (95% CI 68 - 82) | 4 | <5 y | PCV7 | [41] | Israel |
| 86.36% (95% CI 75.78 - 92.32) | 7 | <5 y | PCV7 | [42] | England |
| 98.86% (95% CI 91.91 - 99.84) | 11 | <5 y | PCV7 | [42] | England |
| 96.3% (95% CI 72.97 - 99.49) | 4 | <5 y | PCV13 | [42] | England |
| 71.98% (95% CI 70.15 - 73.71) | 3 | <5 y | PCV13 | [44] | USA |
| 61.46% (95% CI 35.75 - 76.88) | 2 | <5 y | PCV13 | [45] | USA |

| | | | | | |
|---|---|---|---|---|---|
| 94.17% (95% CI 91.48 - 96.01) | 8 | <5 y | PCV7 | [46] | USA |
| 73.92% (95% CI 75.73 – 80.12) | 5 | <5 y | PCV10 | [47] | Kenya |
| 61.45% (95% CI 46.62 – 72.16) | 2 | <5 y | PCV10 | [47] | Kenya |
| 68.84% (95% CI 43.6 - 82.78) | 4 | 2 to <7 y | PCV7 | [43] | USA |
| 49.16% (95% CI 28.24 - 63.98) | 1.5 | >15 y | PCV7 | [48] | Australia |
| 36.57% (95% CI 10.97 - 54.81) | 4.5 | 1-4 y | PCV13 | [49] | Malawi |

**Table S5 – Percent reduction in carriage for several studies.** Mean, 95% CI, age-group, time since vaccination, vaccine, reference and country are presented for each study. See text for mean and CI calculation.

### Literature review of pre-vaccination carriage levels

Table S6 presents a literature review of pre-vaccination carriage levels for different groups of (vaccine) serotypes in different epidemiological contexts. These values are used in figures of the main text comparing results from Blantyre (Malawi) with other countries.

| Carriage in % | Age | Vaccine types | Ref. | Country / Region |
|---|---|---|---|---|
| 53.80% (95% CI 46.02 - 61.44) | <5 y | PCV7 | [39] | USA |
| 27% (95% CI 23 - 32) | <5 y | PCV13 | [17] | Vietnam |
| 67.87% (95% CI 63.44 - 72.07) | <5 y | PCV7 | [35] | Canada |
| 24.7% (95% CI 24.7 - 24.7) | <5 y | PCV7 | [53] | Gambia |
| 53.13% (95% CI 49.40 - 56.84) | <5 y | PCV7 | [33] | Portugal |
| 55.43% (95% CI 50.26 - 60.52) | <5 y | PCV7 | [34] | USA |
| 19.04% (95% CI 10.24 - 30.90) | <5 y | PCV7 | [55] | UK |
| 29.03% (95% CI 21.23 - 37.86) | <5 y | PCV13 | [37] | USA |
| 15.9% (95% CI 13.06 - 18.98) | <5 y | PCV13 | [38] | Italy |
| 46.8% (95% CI 46.8 - 46.8) | <5 y | PCV13 | [53] | Gambia |
| 46.4% (95% CI 41.26 - 51.59) | <5 y | PCV7 | [22] | Nigeria |
| 70.4% (95% CI 65.49 - 74.97) | <5 y | PCV13 | [22] | Nigeria |
| 50.2% (95% CI 44.07 - 56.29) | <5 y | PCV10 | [54] | Nigeria |
| 69% (95% CI 63.12 - 74.45) | <5 y | PCV13 | [54] | Nigeria |

**Table S6 – Pre-vaccination carriage levels for several studies.** Mean, 95% CI, age-group, vaccine types (serotypes of particular PCV vaccine), reference and country are presented for each study.

### Extra results for sensitivity of vaccine impact projections

In Figures S3 and S4 we present extra results on the sensitivity of vaccine impact projections relative to baseline transmission, as an extension to main Figure 5 (and thus using the transmission matrix of Figure S1a. Figure S3 is based on all age-groups under the age of 10 years old as projected by the model (for comparison, the age-groups of main Figure 5 are presented again). Figure S4 presents two age classes for which we found empirical reports on carriage decrease post-vaccination, as an extension to main Figure 5.

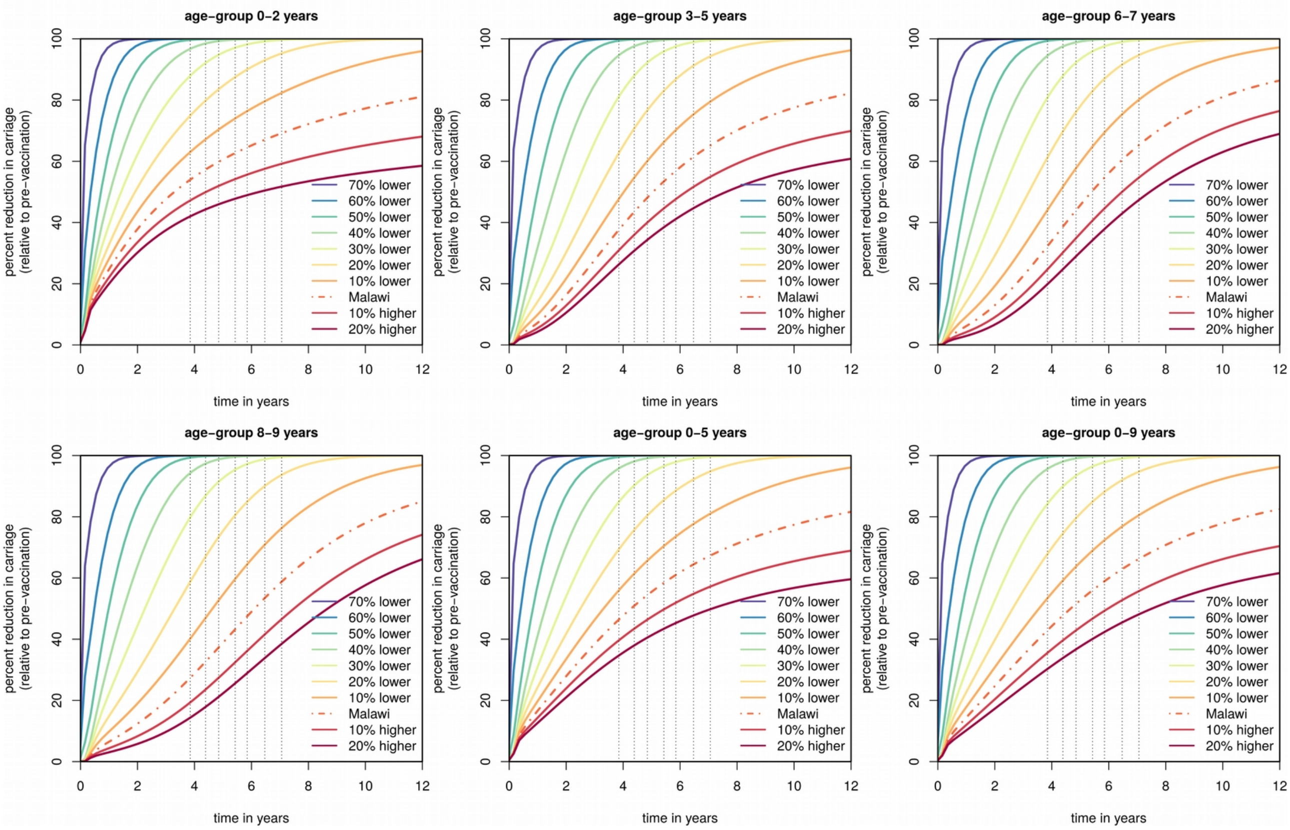

**Figure S3: Extra results for sensitivity of impact projections to baseline transmission (model age-groups under the age of 10 years).** Figure 5 of the main text presents results for age-group 0-5 years of age. This figure presents results for all age-groups. The baseline transmission is varied by considering the 70%, 60%, 50%, 40%, 30%, 20%, 10% lower, and 10%, 20%, 30% higher relative change to the original values estimated for β and θ when fitting the observational study's data (e.g. 40% lower means 0.6*β and 0.6*θ). The impact projections using β and θ of Blantyre (Malawi) are presented with the shaded line (same as in main text). For visual purposes, only the means are shown, obtained from simulations sampling 100,000 parameter values from posteriors.

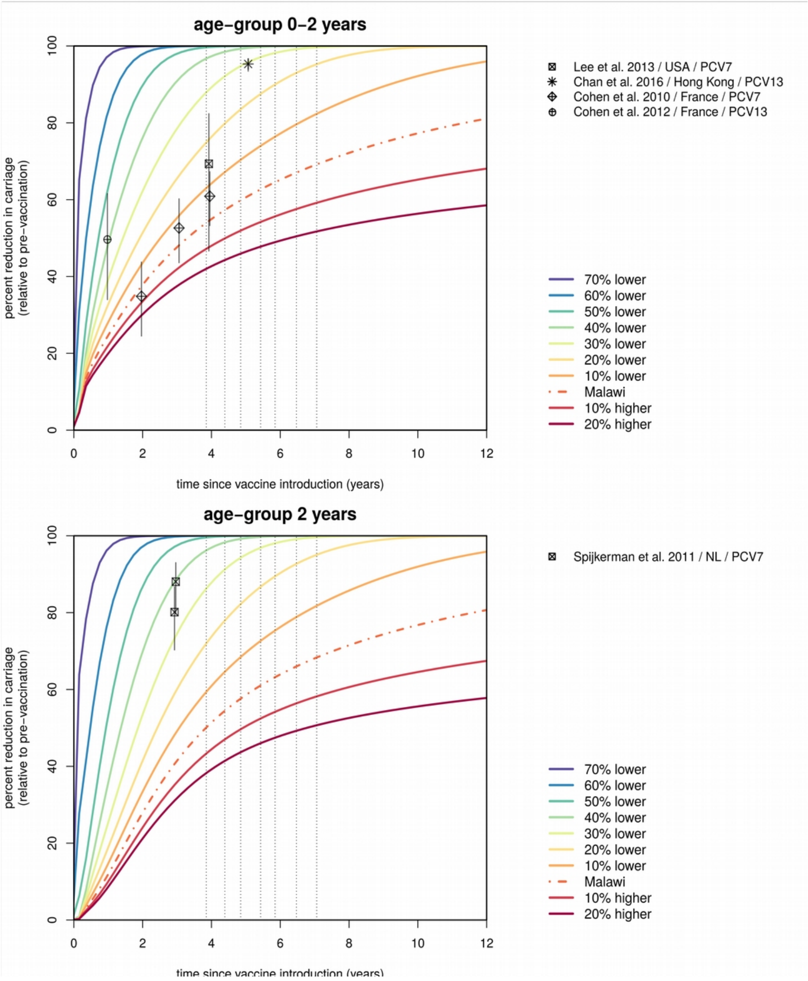

**Figure S4: Extra results for sensitivity of impact projections to baseline transmission (two model age-groups with empirical observations).** Figure 5 of the main text presents results for age-group 0-5 years of age including reported data. This figure presents results for two other age-groups for which reports were available. The baseline transmission is varied by considering the 70%, 60%, 50%, 40%, 30%, 20%, 10% lower, and 10%, 20%, 30% higher relative change to the original values estimated for β and θ when fitting the observational study's data (e.g. 40% lower means 0.6*β and 0.6*θ). The impact projections using β and θ of Blantyre (Malawi) are presented with the shaded line (same as in main text). For visual purposes, only the means are shown, obtained from simulations sampling 100,000 parameter values from posteriors.

### Observational study data

Table S7 keeps the carriage levels observed in the observational surveys and used to fit the model.

| **survey \ age-group** | Vacc. 2 years | Vacc. 3-5 years | Unvacc. 6-7 years | Unvacc. 8-9 years |
|---|---|---|---|---|
| Survey 1 | No data | 0.199 (0.0236) | 0.283 (0.0619) | 0.208 (0.0414) |
| Survey 2 | No data | 0.205 (0.0232) | 0.237 (0.0488) | 0.145 (0.0424) |
| Survey 3 | No data | 0.208 (0.0214) | 0.212 (0.0411) | 0.19 (0.0428) |
| Survey 4 | 0.218 (0.0371) | 0.175 (0.0196) | 0.176 (0.0413) | 0.123 (0.0366) |
| Survey 5 | 0.184 (0.0363) | 0.194 (0.0205) | 0.125 (0.0585) | 0.0851 (0.0407) |
| Survey 6 | 0.191 (0.0337) | 0.15 (0.0183) | 0.132 (0.0465) | 0.112 (0.0353) |
| Survey 7 | 0.151 (0.0319) | 0.167 (0.0207) | 0.2 (0.179) | 0.162 (0.0341) |

**Table S7** – Carriage data from the observational study (mean and standard error)
